# Genetic and Environmental Influences on Pubertal Hormones in Human Hair across Development

**DOI:** 10.1101/152470

**Authors:** Andrew D. Grotzinger, Daniel A. Briley, Laura E. Engelhardt, Frank D. Mann, Megan W. Patterson, Jennifer L. Tackett, Elliot M. Tucker-Drob, K. Paige Harden

## Abstract

**Background:** Puberty is a complex biopsychosocial process that is linked to an array of psychiatric and medical disorders that emerge in adolescence and persist across the lifespan. Although the pubertal process is driven by neuroendocrine changes, few quantitative genetic studies have directly measured puberty-relevant hormones. Hair samples can now be assayed for accumulation of hormones over several months. In contrast to more conventional salivary measures, hair measures are not confounded by diurnal variation or hormonal reactivity.

**Methods:** In an ethnically and socioeconomically diverse sample of 1,286 child and adolescent twins and multiples from 672 unique families, we estimated genetic and environmental influences on concentrations of testosterone, DHEA, and progesterone in hair across the period of 8 to 18 years of age.

**Results:** We identified sex-specific developmental windows of maximal heritability in each hormone. Peak heritability for DHEA occurred at age 9.8 years for males and 10.0 years for females. Peak heritability for testosterone occurred at age 12.5 and 15.2 years for males and females, respectively. Peak heritability for male progesterone occurred at 11.2 years, while the heritability of female progesterone remained uniformly low.

**Conclusion:** This is the first study of genetic influences on pubertal hormone concentrations in human hair and is the largest of the heritability of pubertal hormones in any form. The identification of specific developmental windows when genetic signals for hormones are maximized has critical implications for well-informed models of hormone-behavior associations in childhood and adolescence.

## Genetic and Environmental Influences on Pubertal Hormones in Human Hair across Development

Adolescents who begin puberty at earlier ages are at elevated risk for an array of negative psychiatric and medical outcomes, including depression, eating disorders, substance use, obesity, polycystic ovarian syndrome, and several cancers (1-4). Genome wide association studies support a genetically mediated model of associations between pubertal timing and health (5-7), and have identified specific loci linked to pubertal timing, including genes that are expressed in the ovaries and involved in hormone secretion (8). Puberty involves two distinct sets of hormonal changes — adrenarche and gonadarche — characterized by rising levels of hormones that are themselves associated with health outcomes (9). Earlier pubertal timing, therefore, is an indirect index of early exposure to hormonal risk factors, and a critical step in understanding health associations with pubertal timing involves charting the mechanisms underlying the hormonal bases of individual variation in pubertal development. Toward this end, we estimate quantitative genetic models of genetic and environmental influences on pubertal timing, testosterone, progesterone, and dehydroepiandrosterone (DHEA), and their covariation, in a population representative sample of child and adolescent twins.

### Functions of Pubertal Hormones

Adrenarche and gonadarche are biologically and developmentally dissociable processes characterized by maturation of the hypothalamic-pituitary-adrenal (HPA) and hypothalamicpituitary-gonadal (HPG) axes, respectively (10). Adrenarche begins with the expansion of the inner layer of the adrenal cortex, the zona reticularis, which causes a subsequent increase in DHEA and its sulphate ester (DHEAS; 11). The typical onset of adrenarche is between 6 and 8 years of age, but what causes the initiation of this period is not yet understood (12). DHEA concentrations gradually increase from that point on before peaking in the second decade of life (13). Gonadarche begins approximately two years after adrenarche onset with the increase of gonadal hormones, including testosterone and progesterone. Rising concentrations of DHEA cause the onset of body odor and pubic hair growth, while rising gonadal hormones drive the emergence of secondary sex characteristics in both boys and girls (14; 15).

Pubertal hormones are implicated in a number of socially significant outcomes. DHEA affects immune, metabolic, and central nervous systems (16), and is potentially associated with affective outcomes. For example, 8 to 16 year olds with major depression were characterized by morning DHEA hyposecretion (17), and exogenous administration is reported to increase positive mood in depressed participants (18). These effects may be attributable to the ability of DHEA to function as a GABAA receptor antagonist (19), and bind to both androgen and estrogen receptors (20). Testosterone is associated with adolescent risk taking (21; 22), sexual behavior, substance use (23), and sensation seeking (24). Progesterone is associated with emotional processing (25), and may antagonize androgenic effects (26). In addition, pubertal hormones are capable of affecting the stress response via effects on cortisol synthesis, function, and responsiveness (27-29).

### Genetic Variation in Pubertal Hormones

Although research has made progress in parsing the bidirectional links between pubertal hormones and behavior, the extent to which individual differences in hormones are associated with genetic variation across adolescent development is less clear. Quantitative genetic designs use genetic similarities between relatives (e.g., twins) to disentangle genetic and environmental effects. A handful of previous studies indicate that DHEA levels in adolescence contain a large heritable component in males and females (30; 31). Additive genetic influences on adolescent testosterone have also been identified for both sexes, with three out of five studies indicating higher heritability in males (30; 32-35). Importantly, none of these studies employed measures of long-term hormone levels; only one small study examined age differences in the magnitude of genetic effects; and none have estimated the heritability of progesterone.

Many of the previous studies included participants across the span of adolescence. However, age-specific sex differences in biosynthetic pathways suggest that this may aggregate over dynamic levels of genetic input. For example, DHEA is the primary precursor to testosterone biosynthesis for pre-pubertal children and females across the lifespan, but not for post-pubertal males (36). This likely reflects the fact that testosterone switches from primarily adrenal to gonadal in origin during puberty for males only. In support of the developmental specificity of genetic effects, the heritability of testosterone was found to increase for males (64% to 78%) and decrease for females (70% to 51%) from ages 9 to 12 (34). The key research question, then, is whether we can identify specific periods when hormonal biomarkers are the strongest indicators of an underlying genetic signal (i.e., when heritability is maximized).

### Measurement of Hormones in Hair

Quantitative genetic research on hormone levels is complicated by measurements that are reflective of both state- and trait-specific processes. DHEA, testosterone, and progesterone all display diurnal patterns characterized by peak levels in the morning, followed by a steady decline across the day (37-40). There is also considerable variation in hormones across days, with correlations between testosterone samples taken at the same time of day two days apart estimated at .62 (41). Although multiple salivary or blood samples can be collected to estimate more trait-like hormonal patterns, this can be costly and may decrease participant compliance.

Hair sampling is a recently developed method that provides a measure of long-term, free hormone output using a single, non-invasively collected sample (42). Recent evidence supports the model of hair as a marker of longer-term hormone exposure; in humans, the correlation between a single hair sample and multiple salivary samples assayed for cortisol was shown to linearly increase over a 1-month period as a function of the increased number of collected saliva samples (43).

### Current Study

The current study sought to identify the developmentally-specific and sex-specific genetic and environmental etiologies of hair biomarkers of testosterone, progesterone, and DHEA in a large, population representative sample.

## Methods and Materials

### Participants

Participants were drawn from the Texas Twin Project, a population-based study of school-aged twins (44). Parents of twin families were contacted by mail and invited to participate in an in-lab study. Thirty-one participants were excluded on the basis of reported oral contraception use (see Supplement for hormonal effects) and eight participants were excluded for reported endocrine problems. The current study draws from a final sample of *n* = 1,286 individual twins who provided hair samples from 672 unique families. One hundred and eleven participants had repeat observations, for a total of *i* = 1,397 observation points. The twins were in grades 3 through 12 and ranged in age from 7.80 to 19.47 years (*M* = 12.34, *SD* = 2.77). Two families had repeat triplets with two observations missing at event 2 (5 pairs each), one family had quadruplets (6 pairs), 24 families had triplets (3 pairs), 3 families had triplets with two missing observations (2 pairs), one family had two sets of twins (2 pairs), and 61 pairs were repeat observations (2 pairs), for a total of 798 twin pairs (266 monozygotic [MZ] pairs [108 male, 158 female] and 532 dizygotic [DZ] pairs [112 male, 148 female, and 272 opposite-sex]). Both twins provided hair specimens for 681 pairs out of the 798 twin pairs. Sixty-three percent (63%) of the twins identified as non-Hispanic White, 18.6% identified as Hispanic/Latino, 4.2% identified as African American, and 14.2% identified as another race/ethnicity.

### Measures

#### Zygosity

Opposite-sex twin pairs were classified as DZ. Same-sex twin pair zygosity was assessed using responses to ratings about the twin’s physical similarities (e.g., facial appearance). Two research assistants, high school aged twins, and parents completed the rating. Parents and high school aged twins additionally rated how often they are mistaken for one another. These ratings were entered into a latent class analysis (LCA) that was used to obtain zygosity classifications. LCA is reported to accurately determine zygosity > 99% of the time (45).

#### Pubertal development

Participants rated their pubertal development on the Pubertal Development Scale (46). Boys and girl rated their growth in height, growth in body hair, and skin changes, such as pimples, on a 4-point scale (1=*Not Yet Begun* to 4=*Has Finished Changing*). Only males additionally rated deepening of voice and growth of facial hair, while only girls rated breast development and if they had started menstruating (1 = *No*, 4 *= Yes*). Pubertal status spanned the full range (1 to 4) in both sexes (Figure S1). A total pubertal status score was calculated by averaging across the items. On average, girls (*M* = 2.55, *SD* = 0.98) reported more advanced pubertal status than boys (*M* = 2.07, *SD* = 0.77). Girls also reported their age at menarche (in years). Fifty-five percent (55%) of girls reported having begun menstruating, and among menstruating girls, the average age at menarche was 12.01 years (*SD* = 1.38 years).

#### Hair hormones

Hair samples summing 3 mm in diameter were taken from the posterior vertex of the scalp. Twenty-eight percent (28%) of participants (70% males) had hair shorter than the 3 cm requirement. Based on an average hair growth rate of 1 cm/month (47), the first 3 cm of the hair sample represents a 3-month accumulation of hormones. Samples were stored at room temperature before being shipped to Dr. Clemens Kirschbaum’s laboratory at Technical University Dresden. Participants were still instructed not to use any hair products that are not rinsed out of the hair on the day of the appointment.

Hair samples were analyzed using liquid chromatography-tandem mass spectrometry (LC-MS/MS). Details of this method are described elsewhere (48). The test-retest stability was examined in the 111 repeat participants and indicated high consistency for female progesterone and male DHEA (see Supplement). The inter- and intra-assay coefficients of variation (CVs) have been reported as < 10% for all hair hormones (48), and the lower limit of detection was 0.1 pg/ml for testosterone, DHEA and progesterone. There were 398 (28.5%) samples below the sensitivity threshold for testosterone, 331 (23.7%) samples for progesterone, and 138 (9.9%) samples for DHEA. Binomial regressions indicated that younger participants were significantly more likely to have concentrations below the sensitivity threshold (0 = above threshold, 1 = below threshold) for testosterone (*log odds* = -.07, 95% Confidence Interval (CI) = [-.11, -.02], *p* = .003), progesterone (*log odds* = -.11, 95% CI = [-.16, -.06], *p* < .00 1), and DHEA (*log odds* = - .17, 95% CI = [-.24, -.09], *p* < .001).

### Analyses

Monte Carlo simulations based on the characteristics of the current sample were conducted to determine the best approach to modeling hormone concentrations below the sensitivity threshold. Tobit models—an analytic approach that estimates parameters based on the assumption that values at the lower limit of detection indicate true values occupying an unobserved lower tail of a normal distribution—were most likely to produce 95% confidence intervals that captured the genetic and environmental variance components specified in the generating models (see Supplement). Participants with below threshold values were, therefore, included in all models and data were analyzed using Tobit models estimated with robust weighted least squares (WLSMV). A winsorizing procedure was used to replace extreme hair hormone values by the highest observed score within 3 standard deviations of the sample mean. This involved replacing 11 outliers for hair testosterone, 7 outliers for progesterone, and 24 outliers for DHEA. All three hair hormones were positively skewed, and hormonal measurements were log-transformed to approximate normal distributions in the range of values above-detection limits and then standardized.

As hair assays were performed annually, all models controlled for the year hair samples were analyzed (i.e., batch effects) using a set of nominal variables. Sex was effects coded (-.5 = female, .5 = male), and age was mean centered prior to calculating the quadratic effect. Standard errors and model fit statistics were corrected for non-independence among twins and repeat observations using the complex sampling option (49). Biometric models of the twin correlations were examined using structural equation modeling (SEM) in M*plus* version 7.4. Model fit was evaluated using the root mean square error of approximation (RMSEA; 50) and the comparative fit index (CFI; 51).

## Results

### Effects of Covariates on Hair Hormones

In order to examine the effect of covariates, models were run that included age, sex, race, age^2^, and an age × sex interaction on each hair hormone individually. We first estimated the separate effects of age and sex to yield interpretable main effects (Table S2). DHEA and testosterone were both significantly higher in males. In addition, an age × sex interaction was identified for DHEA and progesterone (Figure 1). Relative to males, female DHEA started lower at age 8, but rose more rapidly across age, such that mean levels were approximately equivalent across sexes by age 18. Progesterone levels were largely stable across age for both sexes, with modest decreases across age observed for males and a slight increase in progesterone levels observed for females. Race was included as a covariate for hormones and puberty in remaining models, and the effect of sex was taken into account by allowing for separately estimated means across males and females. Sex-specific effects of age and age^2^ were also estimated unless otherwise noted.

**Figure 1.**
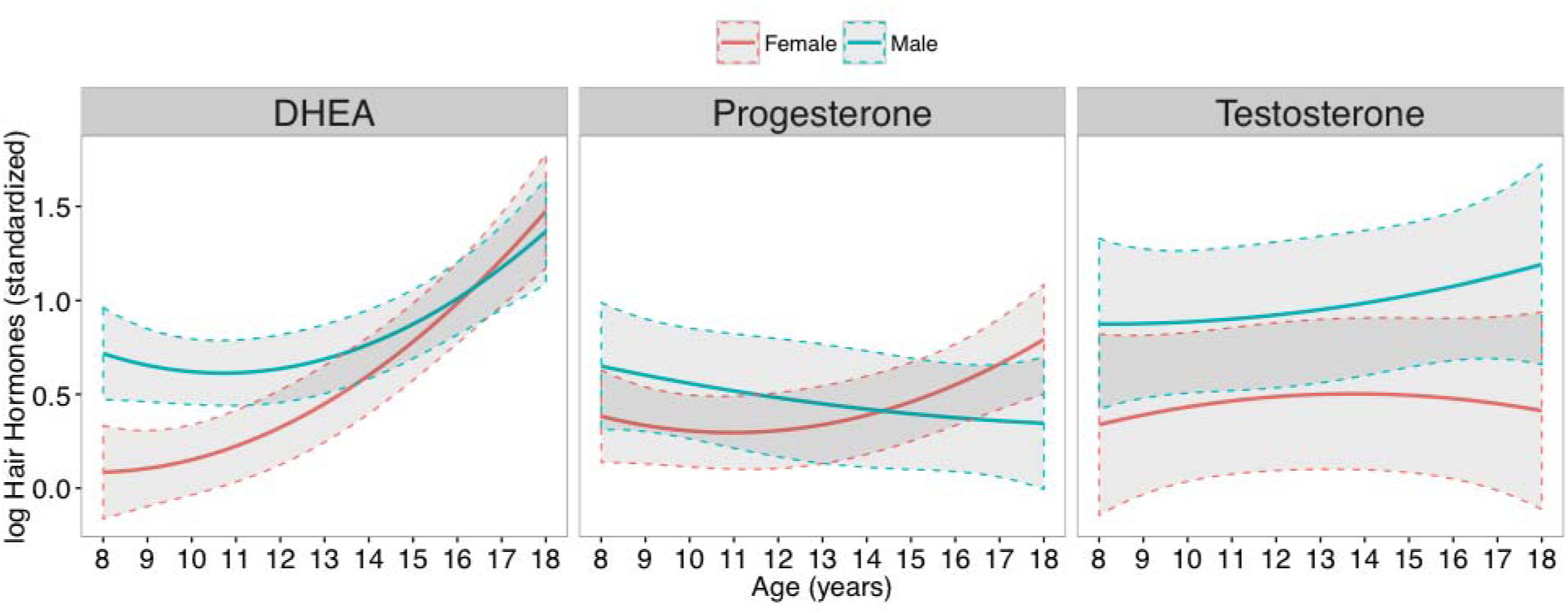
Quadratic age trends for hair hormones for males and females. Hair hormones were winsorized, log transformed, and standardized across sex. Dashed lines indicate 95% confidence intervals.

### Phenotypic Associations between Hair Hormones and Puberty

#### Model specification

We next examined the phenotypic relations among hair hormone levels, age, and self-reported pubertal development in each sex using multiple group models. The residuals between the hormones were freely correlated for all models. The first model tested the direct association between hormones and pubertal development, without controlling for age (Model A). The second model (Model B) included the sex-specific linear and quadratic effects of age on hormone levels and pubertal development, and tested for mediation by hair hormones of age effects on puberty. Notably, removing variance in pubertal status due to age changes the meaning of the variable; it now represents pubertal *timing*, i.e., the extent to which a child/adolescent is more or less developed than his or her same-age peers.

Finally, we used nonparametric local structural equation models (LOSEM) to examine the associations between hair hormones and pubertal timing across age as a continuous moderator (Model C; 52). LOSEM functions by using a weighting kernel and bandwidth that assigns a higher weight to observations in closer proximity to the focal value of the continuous moderator. Multiple structural equation models were estimated that differed only with respect to the assigned focal value of age, which ranged from 8 to 18 years at intervals of .10. Age was excluded as a covariate as the LOSEM models restrict the variance in age, resulting in age effects estimated near 0. Pubertal timing was, therefore, calculated by residualizing pubertal status for sex-specific linear and quadratic effects of age prior to fitting LOSEMs to the data.

#### Model results

Results from all phenotypic models of puberty are depicted in Figure 2. In Model A, higher DHEA was associated with more advanced pubertal development in both sexes, while progesterone was associated with self-reported puberty only in females, and testosterone with puberty only in males. Hair hormones accounted for 8.2% of the variance in pubertal development for females, and 4.6% of the variance for males. In Model B, which added the direct effects of age, older females reported more advanced pubertal development and had higher levels of progesterone and DHEA. For males and females, hair hormones were not predictive of pubertal timing, and the total indirect effect of age and age^2^ on puberty via hormones was negligible ( all *β*s < .01.)

**Figure 2.**
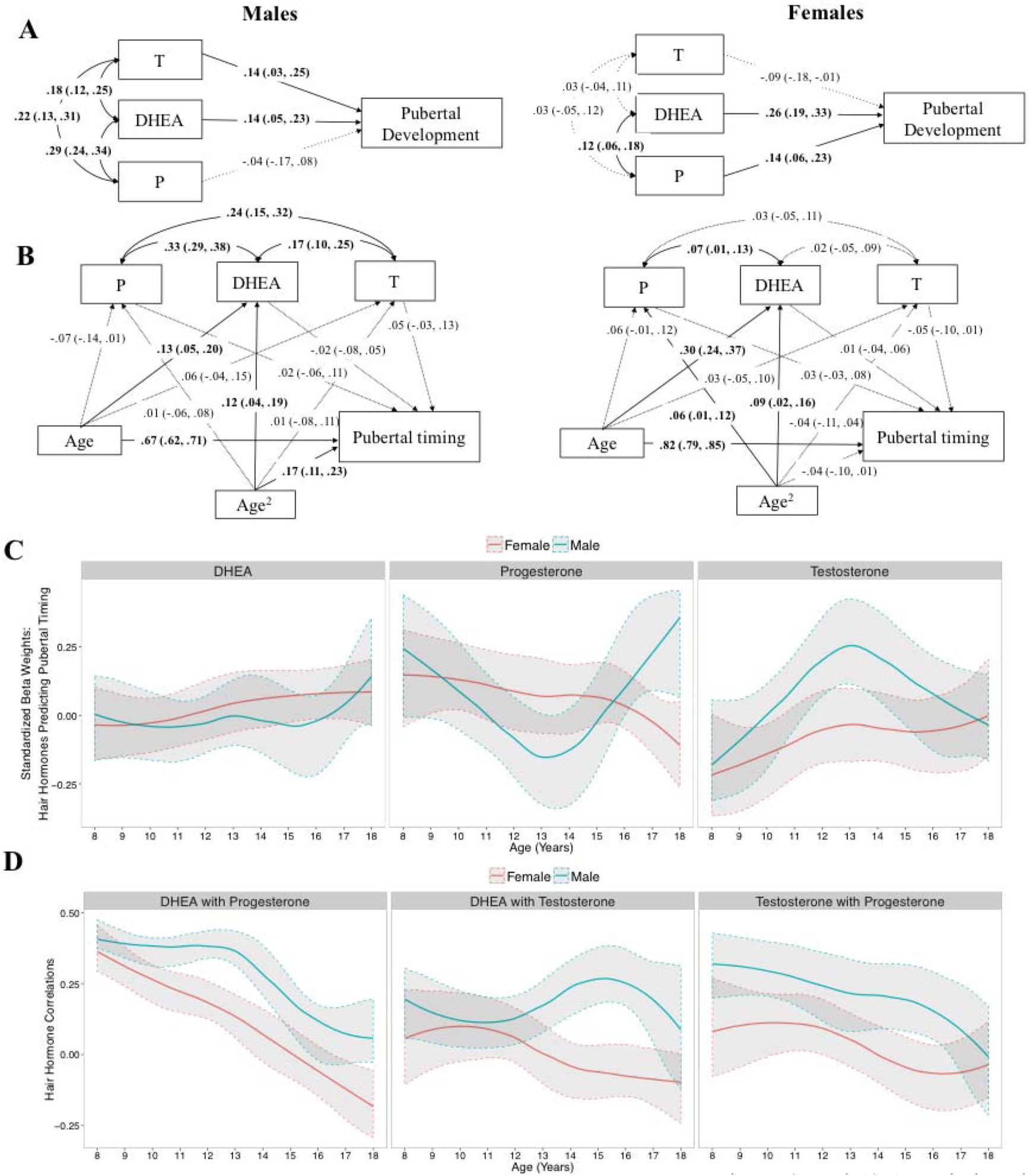
Phenotypic associations between puberty and hair hormones across age and sex. (Panel A) Associations between hair hormones and pubertal status in males (left panel) and females (right panel). (Panel B) Associations between hair hormones and pubert timing (i.e., pubertal status controlling for the effects of age). Standardized parameter estimates with 95% confidence intervals are provided in parentheses. Parameters significantly different from zero at *p* < .05 are illustrated with solid lines; parameters not significantly different from zero at *p* < .05 are illustrated with dashed lines. (Panel C) Associations between hair hormones and puberta timing as a function of age estimated using LOSEM. (Panel D) Correlations among hair hormones as a function of age estimated using LOSEM. Gray bands indicate 95% confidence intervals. All models included the effects of race on puberty and hormones and analytic batch on hormones (not shown). T = testosterone. P = progesterone. DHEA = dehydroepiandrosterone.

LOSEM results indicated that the association between DHEA and pubertal timing increased slightly across age for both sexes. Progesterone levels were moderately associated with female pubertal timing, though the association decreased starting at ~15 years of age. Male progesterone levels evinced a U-shaped relationship with pubertal timing, predicting the most variance at ages 8 and 18, and the least variance at age 13. This was in contrast to male testosterone levels that showed an *inverted* U-shaped pattern, with a rise in the association from age 8 that peaked at age 13, followed by a steady decrease thereon. Female testosterone levels were modestly associated with pubertal timing across the full age range. Residual correlations among the hair hormones from the same LOSEM models are depicted in Figure 2. Correlations between progesterone and both DHEA and testosterone decreased across age for males and females. The correlation between DHEA and testosterone decreased across age for females, but evinced a modest increase from ages 12 to 15 for males.

### Univariate Twin Models

#### Twin correlations

Cross-twin correlations are summarized in Table 1. The correlations among MZ twins’ hormone levels were generally high, suggesting low levels of non-shared environmental input. The difference between MZ and DZ correlations was higher for males than females for DHEA and testosterone, indicating larger genetic influences in males.

**Table 1.** Cross-twin correlations for hair hormones, pubertal status and pubertal timing

|  | Monozygotic |  | Dizygotic |  |  |
| --- | --- | --- | --- | --- | --- |
|  | Male | Female | Male | Female | Opposite-sex |
| Pubertal Status | .69 (.58, .79) | .78 (.69, .88) | .65 (.55, .76) | .74 (.64, .83) | .64 (.57, .72) |
| Pubertal Timing | .28 (.05, .51) | .40 (.22, .57) | .30 (.16, .44) | .23 (.04, .42) | .13 (-.04, .30) |
| DHEA | .78 (.64, .92) | .58 (.45, .71) | .28 (.14, .42) | .66 (.50, .82) | .37 (.27, .47) |
| Progesterone | .67 (.55, .78) | .62 (.54, .70) | .91 (.77, 1.05) | .91 (.69, 1.13) | .46 (.36, .56) |
| Testosterone | .94 (.74, 1.14) | .62 (.50, .73) | .65 (.48, .82) | .42 (.29, .55) | .49 (.38, .60) |

*Note.* All models included race as a covariate and allowed for sex-specific means and residual variances in outcomes. Hormonal outcomes were regressed on analytic batch and all outcomes except pubertal status were regressed on sex-specific effects of age and age-squared. 95% confidence intervals are given in parentheses.

#### Twin model specification

Quantitative genetic models were fit to the data to determine variance attributable to additive genetic (*A*), shared environmental (*C*) and non-shared environmental factors, including error variance, unique to each twin (*E*). The *ACE* components were standardized. As MZ and DZ pairs share approximately 100% and 50% of their genes, respectively, the *A* factor is fixed to correlate at 1.0 for MZ pairs and 0.5 for DZ pairs. Correlations between the *C* factors are fixed to 1.0 in all twin pairs. Each hormone, pubertal status, and pubertal timing were examined in a series of five univariate models. All five models provided good fit to the data (CFIs ≥ . 987, RMSEAs ≤ .028

#### Model results

An initial test of scalar sex differences—that is, differences in the age-independent variance that could be accounted for by *ACE* factors—indicated that there was significantly more variance in female DHEA and significantly less variance in female progesterone and pubertal status. For this reason, we compare proportions of variance, as opposed to unstandardized variance, accounted for by *ACE* factors across the sexes (summarized in Table 2; displayed in Figure 3). DHEA levels were significantly more heritable in males (63%) relative to females (10%). Female testosterone was influenced by a significantly larger non-shared environmental effect (38% vs. 7% in males). Testosterone was primarily heritable in males (55%) and females (44%). Variation in pubertal status and progesterone was largely due to shared environment in both sexes, and variance in pubertal timing was predominantly attributable to non-shared environmental influences.

**Table 2.** Proportion of Variance and Sex Differences from Univariate Models

|  | Males |  |  | Females |  |  | Males - Females |  |  |  |
| --- | --- | --- | --- | --- | --- | --- | --- | --- | --- | --- |
| | $h^2$ | $c^2$ | $e^2$ | $h^2$ | $c^2$ | $e^2$ | $\Delta h^2$ | $\Delta c^2$ | $\Delta e^2$ | $\Delta \text{phenotypic variance}$ |
| Pubertal Status | .16 (.17) | .53 (.13)*** | .31 (.06)*** | .09 (.21) | .69 (.17)*** | .22 (.06)*** | .07 (.33) | -.16 (.26) | .09 (.09) | -.41 (.06)*** |
| Pubertal Timing | .05 (.34) | .40 (.21)** | .55 (.22)* | .36 (.21) | .06 (.19) | .58 (.06)*** | -.31 (.55) | .34 (.38) | -.03 (.24) | -.03 (.03) |
| DHEA | .63 (.14)** | .09 (.10) | .28 (.06)*** | .10 (.21) | .54 (.17)** | .36 (.06)*** | .53 (.14)*** | -.45 (.11)*** | -.08 (.07) | -.34 (.06)*** |
| Progesterone | .04 (.18) | .74 (.14)*** | .22 (.05)*** | .26 (.24) | .40 (.22) | .34 (.04)*** | -.22 (.41) | .34 (.36) | -.12 (.07) | .13 (.05)** |
| Testosterone | .55 (.30) | .38 (.22) | .07 (.10) | .44 (.17)* | .18 (.12) | .38 (.06)*** | .11 (.35) | .20 (.27) | -.31 (.11)** | .10 (.08) |

*Note.* All models included race and sex-specific effects of age and age-squared as covariates and allowed for sex-specific means and
residual variances in outcomes, with the exception of excluding age effects from the pubertal status model. Hormonal outcomes were additionally regressed on analytic batch. The last column on the right reports the difference in the residual variance across sexes, and the adjacent three columns report differences in proportions of variance attributable to *ACE* factors. Standard errors are given in parentheses.
\*\*\* significantly different than zero at $p < .001$ ; \*\* $p < .01$ ; \* $p < .05$

**Figure 3.**
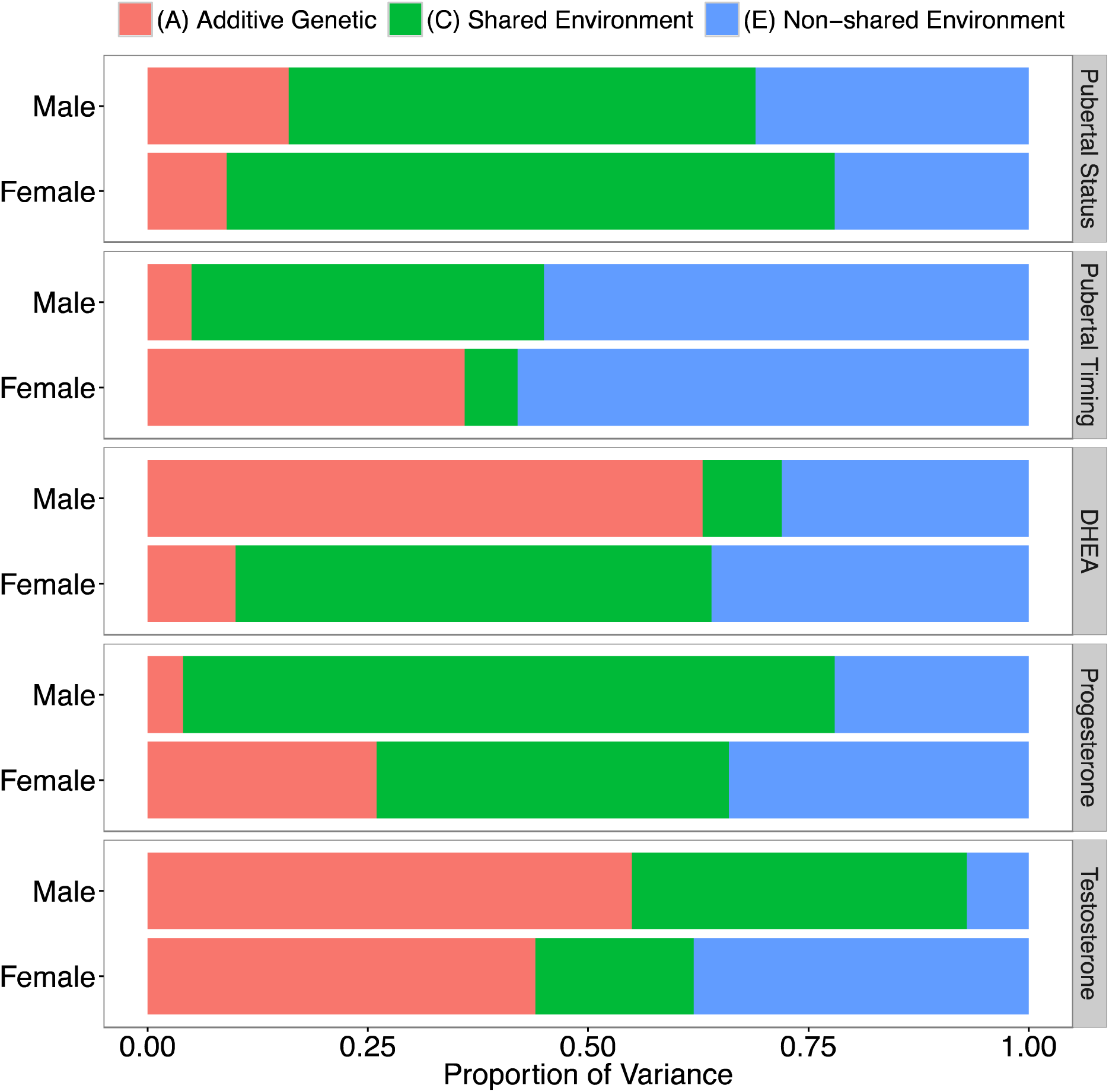
Proportion of variance in each outcome explained by genetic (*A*) and environmental (*C* or *E*) factors from univariate models that allowed for sex differences in all *ACE* paths.

Univariate LOSEMs were then used to examine changes in sex-specific *ACE* estimates across age. Race was excluded as a covariate for biometric LOSEM models as there were limited numbers of African American twins for certain age groups. Unstandardized variance by sex is depicted in Figure 4, and proportions of variance are shown in Figure S2. Estimates for pubertal timing indicated that variability was due to a combination of genes and environment, but that the highest levels of heritability were observed at age 11.6 for females and age 13.2 for males. Male progesterone contained a moderate heritable component from ages 8 to 14 with a peak of 65% heritability at age 11.2 years. Conversely, variation in female progesterone was almost entirely attributable to environmental influences across the age range.

**Figure 4.**
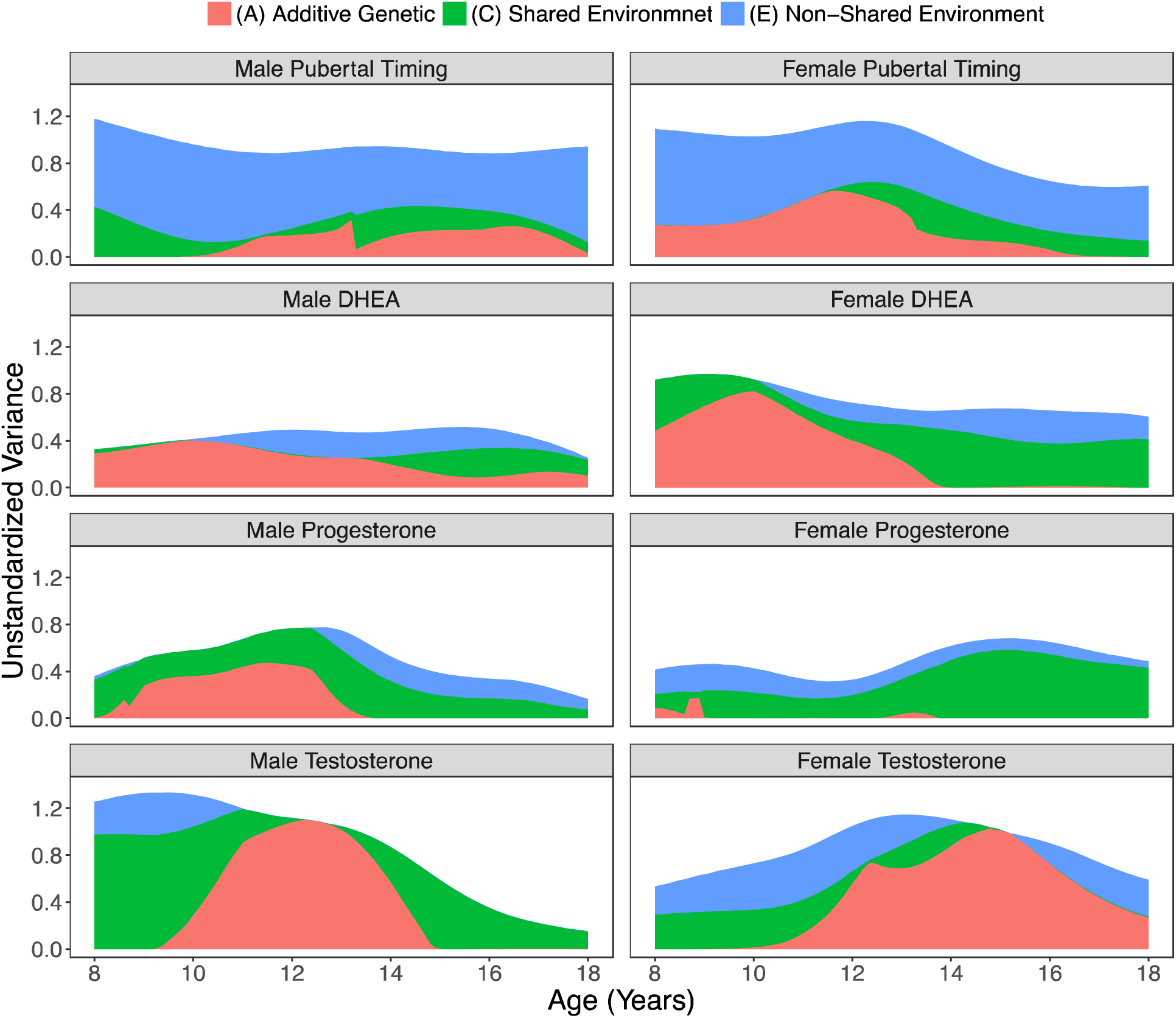
Unstandardized levels of variance explained by *ACE* components from univariate models estimated using LOSEM across age.

Additive genetic influences on male DHEA steadily declined across age, reaching levels of ~20% by age 16. The effect of genes on female DHEA peaked at age 10, followed by a drop to near 0 by age 14. It is of note that DHEA was highly heritable at its peak in both sexes (99% at age 9.8 in males, 89% at age 10 in females). Male testosterone evinced a gradual rise in additive genetic influences starting at age 9, followed by a peak of 100% heritability at age 12.5, and subsequent decline to 0% heritability by age 15. Female testosterone showed a similar, but later, pattern starting at age 10, and reaching 100% heritability at age 15.2.

### Bivariate Twin Models

A series of six bivariate correlated factors models were then used to examine genetic and environmental sources of covariation between DHEA, progesterone, testosterone, and pubertal status. Bivariate associations with pubertal timing are not reported as the phenotypic correlations were estimated near 0 in both sexes, and twin models indicated this was not due to countervailing genetic and environmental input. Table 3 reports the correlations between additive genetic (*r*A), shared environmental (*r*C) and non-shared environmental factors (*r*E), as well as the contributions to phenotypic correlations using the within-sex standardized *ACE* path estimates.

**Table 3.**
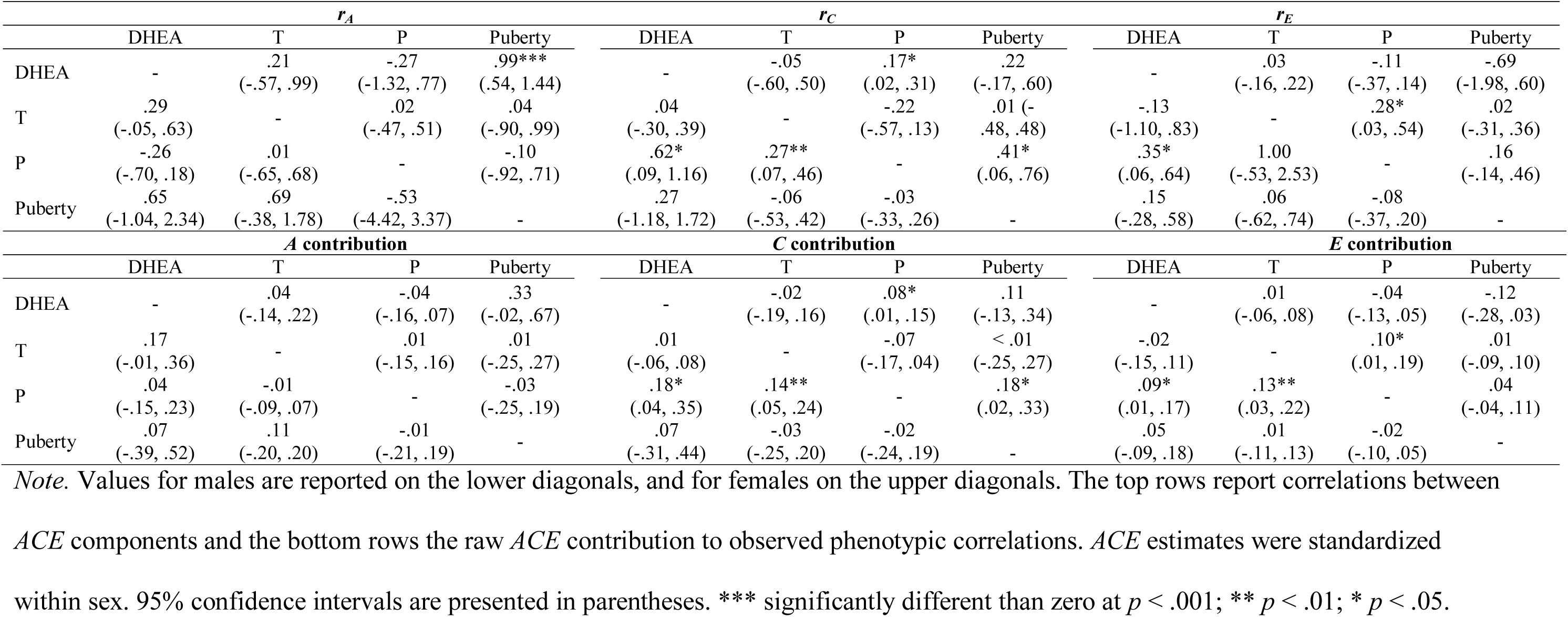
*ACE* correlations and contributions to phenotypic correlations

Additive genetic predictors of DHEA and pubertal status were highly correlated in females (*r_A_* = .99), and the genetic correlation between DHEA and testosterone approached significance for males (*r_A_* = .29, *p* = .10). There was a significant correlation with progesterone via shared environmental pathways for male testosterone (.14), female pubertal status (.18), and DHEA in males (.18) and females (.08). In addition, there were significant correlations with progesterone attributable to non-shared environmental effects for male DHEA (.09), and testosterone in males (.13) and females (.10).

Bivariate results revealed five significant sex differences. The shared environmental correlation between testosterone and progesterone was significantly higher in males (Δ *=* .49, *SE* = .19, *p* = .008), as was the corresponding contribution to the phenotypic correlation (Δ *=* .2 1, *SE* = .07, *p* = .001). Conversely, the shared environmental correlation between progesterone and pubertal status was significantly higher in females (Δ *=* -.44, *SE* = .17, *p* = .010). The correlation between non-shared environmental predictors of DHEA and progesterone (Δ *=* .46, *SE* = .20, *p* = .0 18), and the contribution to the observed correlation (Δ *=* .13, *SE* = .06, *p* = .03 6), was higher in males.

## Discussion

The current paper presents findings from the largest twin study of pubertal hormones to date, and the first genetically informative study of a novel measure – concentrations of puberty-relevant hormones in hair. On average, genetic influences on DHEA were higher in males, and testosterone was highly heritable in both sexes. Variation in progesterone and both pubertal status and timing were largely attributable to environmental influences.

Age-specific analysis, however, indicated that these estimates aggregate over dramatic developmental changes. In both sexes, the heritability of DHEA peaked at age 10 and subsequently declined. For testosterone, heritability was near zero at age 8, increasing to 100% at age 12.5 years in males and 15.2 years in females. This suggests that genetic effects on pubertal hormones become heightened during developmental windows. Point estimates of heritability for pubertal hormones in age-heterogeneous child and adolescent samples are likely to be of limited value.

The current findings regarding the developmental genetics of pubertal hormones may offer insights into the mechanisms underlying behavioral development and psychiatric morbidity. Animal models indicate that pubertal hormones structurally reorganize the brain, particularly in subcortical areas and reward-related systems (reviewed in 53). For example, rats castrated before puberty have also been shown to have higher levels of androgen receptors in the amygdala than rats castrated post-puberty (54), indicating that hormone levels fine-tune their own receptor systems within a specific pubertal window. An emerging literature suggests hormone-driven reorganization of the brain also applies to puberty in humans (55). For example, testosterone levels in adolescence are associated with decreased gray matter volumes in the hippocampus and increased amygdala volume in both sexes (56). DHEA levels have also been associated with increased cortical thickness specifically from ages 4-13 (57). In addition, inverted U-shaped patterns for human gray matter volumes in the frontal and parietal lobes are strikingly similar to the genetic trends observed in the current study (58). As variation in pubertal hormones appears to contain a strong heritable component during neurobiologically sensitive periods, it is critical for future research to identify what causes these genetic signals to come online.

Adolescence is a unique developmental period that involves sweeping biological, psychological, and social changes that reverberate through the rest of the lifespan. Indeed, ~75% of adults with mental disorders have been found to receive diagnoses before age 18, suggesting prevention efforts would do well to target processes that emerge prior to adulthood (59). Pubertal hormones may be a key component of various etiological pathways related to psychiatric risk and onset, though likely through nuanced, developmentally specific mechanisms that are not yet fully understood. The current study adds to a presently sparse literature on genetic variation in hormones, and is the first to identify developmental periods of heightened genetic influences on pubertal hormones. Future work may benefit from testing whether these periods of heightened heritability mark periods of reorganizational effects on neural systems that canalize trajectories of psychiatric symptomology.

## Acknowledgements

This research was supported by National Institutes of Health (NIH) grants R01HD083613, R21HD081437, and R21AA023322. LEE was supported by a National Science Foundation Graduate Research Fellowship. The Population Research Center at the University of Texas at Austin is supported by NIH grant R24HD042849.

